# LIN37-DREAM Prevents DNA End Resection and Homologous Recombination at DNA Double Strand Breaks in Quiescent Cells

**DOI:** 10.1101/2021.04.21.440786

**Authors:** Bo-Ruei Chen, Yinan Wang, Anthony T. Tubbs, Dali Zong, Faith C. Fowler, Nicholas Zolnerowich, Wei Wu, Amelia Bennett, Chun-Chin Chen, Wendy Feng, Andre Nussenzweig, Jessica K. Tyler, Barry P. Sleckman

**Affiliations:** Division of Hematology and Oncology, University of Alabama at Birmingham, Birmingham, AL 35233; O’Neal Comprehensive Cancer Center, University of Alabama at Birmingham, Birmingham, AL 35233; Department of Pathology and Laboratory Medicine, Weill Cornell Medicine, New York, NY 10065; Laboratory of Genome Integrity, National Cancer Institute, Bethesda, MD 20892

## Abstract

DNA double strand break (DSB) repair by homologous recombination (HR) is thought to be restricted to the S- and G_2_- phases of the cell cycle in part due to 53BP1 antagonizing DNA end resection in G_1_-phase and non-cycling quiescent (G_0_) cells. Here, we show that LIN37, a component of the DREAM transcriptional repressor, functions in a 53BP1-independent manner to prevent DNA end resection and HR in G_0_ cells. Loss of LIN37 leads to expression of HR proteins, including BRCA1, BRCA2, PALB2 and RAD51, and DNA end resection in G_0_ cells even in the presence of 53BP1. In contrast to 53BP1-deficiency, DNA end resection in LIN37-deficient G_0_ cells depends on BRCA1 and leads to RAD51 filament formation and HR. LIN37 is not required to protect DNA ends in cycling cells at G_1_-phase. Thus, LIN37 regulates a novel 53BP1-independent cell phase-specific DNA end protection pathway that functions uniquely in quiescent cells.

## Introduction

DNA double strand breaks (DSBs) are repaired by two main pathways, non-homologous end joining (NHEJ) and homologous recombination (HR) (Prakash et al. 2015, Chang et al. 2017). HR functions in the S- and G_2_-phases of the cell cycle using the sister chromatid as a template for precise homology-directed repair of DSBs. In contrast, NHEJ functions in all phases of the cell cycle to rejoin broken DNA ends and is the only pathway of DSB repair in G_1_-phase cells and in non-cycling cells that have exited the cell cycle and are quiescent in G_0_, which comprise the majority of cells in the human body (Chang et al. 2017). The initiation of HR requires DNA end resection to generate extended single stranded DNA (ssDNA) overhangs that are coated by the trimeric ssDNA binding protein complex RPA1/2/3 (hereafter referred to as RPA) (Ciccia and Elledge 2010). RPA is subsequently replaced by RAD51, and the RAD51 nucleofilament mediates a search for a homologous template usually within the sister chromatid to enable the completion of HR-mediated DNA DSB repair (Mimitou and Symington 2009, Wyman, Ristic, and Kanaar 2004). In contrast, NHEJ works best on DNA ends with minimal ssDNA overhangs, necessitating that DNA end resection be limited in G_1_-phase and G_0_ cells (Chang et al. 2017, Symington and Gautier 2011). Extensive resection and ssDNA generation at broken DNA ends in cells at G_0_/G_1_ would antagonize NHEJ-mediated DSB repair and promote aberrant homology-mediated repair, due to the absence of sister chromatids, forming chromosomal translocations and deletions leading to genome instability (Ciccia and Elledge 2010). Therefore, the generation of ssDNA at broken DNA ends is the critical decision point for whether a DSB will be repaired by NHEJ and HR. As such, highly regulated processes that control DNA end resection are critical for ensuring appropriate choice of DSB repair pathways in all cells.

During HR, BRCA1 initiates the resection of broken DNA ends with the CTIP and MRE11 nucleases generating short ssDNA tracts that are extended by the nucleases such as EXO1 and DNA2-BLM (Symington and Gautier 2011). Other proteins, including Fanconi Anemia proteins such as FANCD2, are also involved in regulating DNA end resection (Unno et al. 2014, Murina et al. 2014, Cai et al. 2020). In cells where NHEJ must repair DSBs, the extensive resection of DNA ends must be prevented and 53BP1 and its downstream effectors RIF1 and the shieldin complex antagonize DNA end resection in these cells (Setiaputra and Durocher 2019, Mirman and de Lange 2020, Bunting et al. 2010). 53BP1 and shieldin may protect DNA ends by inhibiting the recruitment or activity of pro-resection proteins and also by promoting DNA synthesis at resected DNA ends to “fill in” the ssDNA gap generated by resection (Mirman and de Lange 2020, Setiaputra and Durocher 2019).

Whether pathways exist in addition to 53BP1 and RIF1/shieldin that protect DNA ends and promote genome integrity by preventing aberrant homology-mediated DSB repair is unclear. Here we develop an unbiased whole genome CRISPR/Cas9 screening approach based on assaying RPA loading at DSBs to identify genes encoding proteins that prevent DNA end resection and ssDNA generation in cells with 2N DNA content, which will be in either G_0_ or G_1_. This approach identified gRNAs targeting genes encoding proteins known to protect DNA ends such as 53BP1 and RIF1. However, this screen also identified LIN37, a protein not known to function in DNA end processing and DSB repair. LIN37 is a component of the DREAM (dimerization partner, RB-like, E2F and multi-vulval) transcriptional repressor complex (Litovchick et al. 2007, Sadasivam and DeCaprio 2013, Mages et al. 2017). Here we show that LIN37 functions in a 53BP1-independent manner to protect DNA ends from resection exclusively in quiescent G_0_ cells. Moreover, while loss of either 53BP1 or LIN37 leads to loading of RPA at DSBs in G_0_ cells, loss of LIN37 further leads to subsequent steps of HR including RAD51 loading and homology-mediated DSB repair.

## Results

### Identification of novel factors that regulate DNA end resection

Murine pre-B cells transformed with the Abelson murine leukemia virus kinase, hereafter referred to as abl pre-B cells, are rapidly cycling cells (Rosenberg, Baltimore, and Scher 1975). When treated with the abl kinase inhibitor, imatinib, abl pre-B cells stop cycling with a predominantly 2N DNA content indicative of them being in G_0_/G_1_ (Bredemeyer et al. 2006, Muljo and Schlissel 2003). Imatinib treated abl pre-B cells that ectopically express Bcl2 survive in culture for several days and we refer to these cells as non-cycling abl pre-B cells (Bredemeyer et al. 2006).

DNA DSBs in G_0_/G_1_ cells are repaired by NHEJ, which requires DNA Ligase 4 to join the broken DNA ends (Chang et al. 2017). Consequently, DNA DSBs generated in non-cycling DNA Ligase 4-deficient (*Lig4^-/-^*) abl pre-B cells persist un-repaired (Helmink et al. 2011). These persistent DSBs are protected from nucleolytic activity and exhibit minimal DNA end resection (Tubbs et al. 2014, Dorsett et al. 2014, Helmink et al. 2011). However, loss of H2AX or 53BP1 in abl pre-B cells leads to extensive DNA end resection and the generation of ssDNA overhangs (Tubbs et al. 2014, Dorsett et al. 2014, Helmink et al. 2011). ssDNA at resected DNA ends binds to RPA, which can be assayed by flow cytometry after detergent extraction of soluble RPA (Forment, Walker, and Jackson 2012). Indeed, as compared to non-cycling *Lig4^-/-^* abl pre-B cells, the generation of DSBs by ionizing radiation (IR) in non-cycling *Lig4^-/-^:53bp1^-/-^* abl pre-B cells led to robust chromatin-associated RPA as assayed by flow cytometry (Figure 1A). This assay detects more extensive DNA end resection in the absence of 53BP1 and thus serves as the basis for a screen to identify additional proteins that normally function to prevent extensive DNA end resection in G_0_/G_1_ cells.

**Figure 1.**
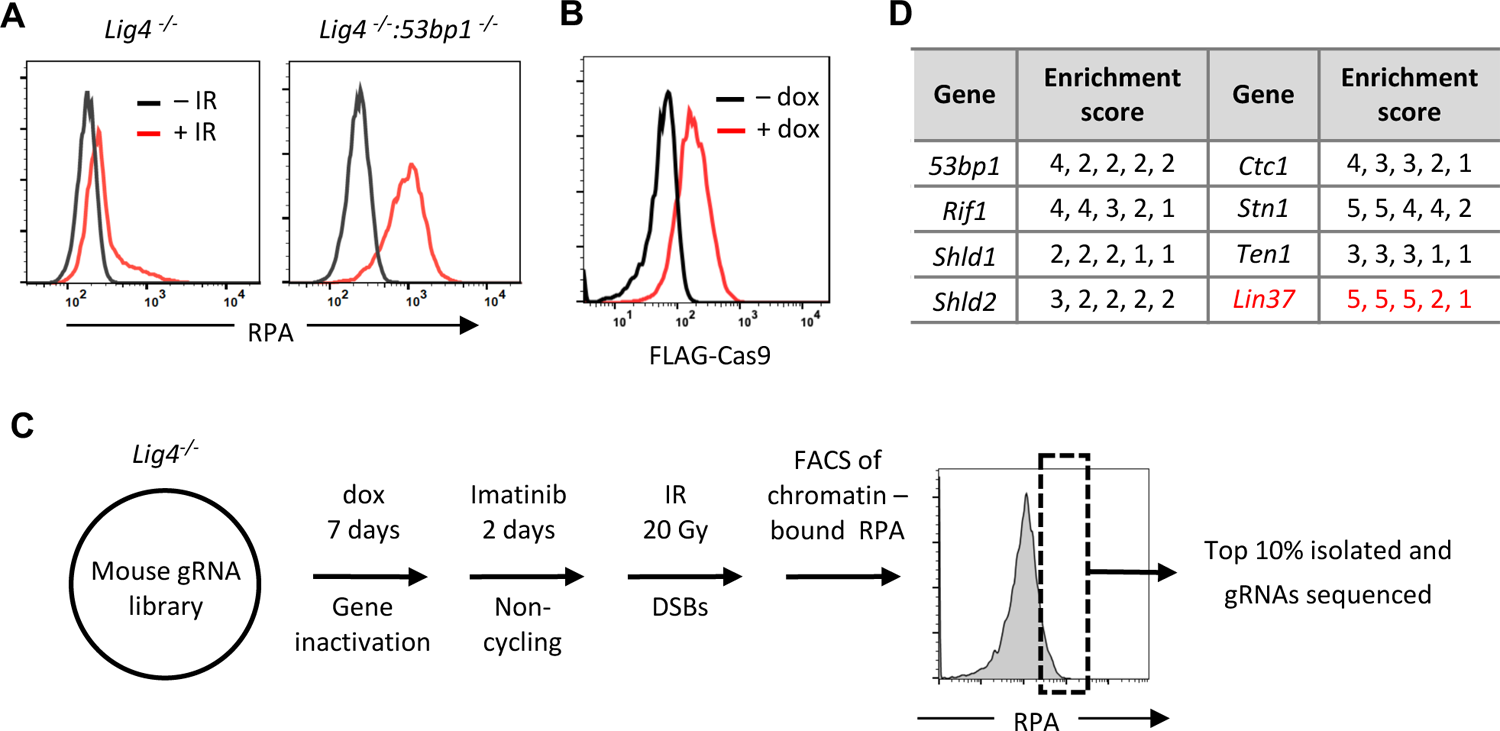
An unbiased genome-scale gRNA screen for novel DNA end protection factors (A) Flow cytometric analysis of chromatin-bound RPA before and after IR of non-cycling *Lig4*^-/-^ and *Lig4*^-/-^:*53bp1*^-/-^ abl pre-B cells. (B) Flow cytometric analysis of FLAG-Cas9 in *Lig4*^-/-^ cells with (+dox) and without (-dox) doxycycline to induce expression of FLAG-Cas9. (C) Schematic diagram of the genome-scale guide RNA screen for genes preventing DNA end resection in non-cycling *Lig4*^-/-^ abl pre-B cells. (D) Enrichment score (fold enrichment) of individual gRNAs to a subset of genes identified in the RPA high population.

To facilitate our screen for gRNAs that increase association of RPA with DSBs, we introduced a lentivirus containing a FLAG-Cas9 cDNA under control of a doxycycline-inducible promoter into *Lig4*^-/-^ abl pre-B cells. Immunodetection of the FLAG epitope allowed flow cytometric assessment of FLAG-Cas9 protein expression upon doxycycline treatment, showing that FLAG-Cas9 can be induced in all the cells in the population (Figure 1B). A lentiviral mouse guide RNA (gRNA) library containing approximately 90,000 gRNAs to 18,000 mouse genes was introduced into these cells followed by doxycycline treatment for seven days to promote Cas9-gRNA-mediated gene inactivation (Figure 1C) (Tzelepis et al. 2016). After 2 days of imatinib treatment to render these cells non-cycling, the cells were subjected to irradiation (IR) and those with the highest level (top 10%) of chromatin-bound RPA were isolated by flow cytometric cell sorting (Figure 1C). The gRNAs from this “RPA-high” cell population were sequenced, and the frequency of individual gRNAs in the RPA-high population relative to those in the “RPA-low” population was determined.

Multiple gRNAs to genes encoding proteins known to prevent DNA end resection such as 53BP1, RIF1 and shieldin subunits were enriched in the RPA-high cell population, demonstrating the veracity of this screening approach (Figure 1D and Table S1). In addition, several gRNAs to *Lin37*, a gene encoding the LIN37 subunit of the MuvB complex were also highly enriched in RPA-high cells (Figure 1D and Table S1) (Litovchick et al. 2007). When bound to the pocket proteins p130/p107, and E2F proteins E2F4/5 and DP, the MuvB complex forms the DREAM transcription repressor complex, which functions to repress genes that promote entry into the cell cycle (Litovchick et al. 2007, Sadasivam and DeCaprio 2013, Mages et al. 2017).

### RPA accumulation at DNA DSBs in non-cycling LIN37-deficient cells

To validate the hit from the gRNA screen, we generated DNA Ligase 4-deficient abl pre-B cells that were deficient in LIN37 (*Lig4^-/-^:Lin37^-/-^*) (Figure 2A). Similar to 53BP1-deficient abl pre-B cells, LIN37-deficienct cells also exhibited increased RPA association with DSBs in chromatin after IR, with most of the cells lacking LIN37 having even more chromatin bound RPA after IR than cells lacking 53BP1 (Figure 2B). This is not a consequence of more DNA damage occurring in the absence of LIN37, because all the cells had similar levels of γH2AX indicative of similar DNA DSB generation in response to IR (Figure 2B). Analyses of non-cycling wild type (*WT*), *53bp1^-/-^* and *Lin37^-/-^* abl pre-B cells that expressed Ligase 4 yielded similar results, indicating that RPA binding at DSBs in 53BP1 and LIN37 deficient abl pre-B cells does not depend on DNA Ligase 4 deficiency (Figure S1).

**Figure 2.**
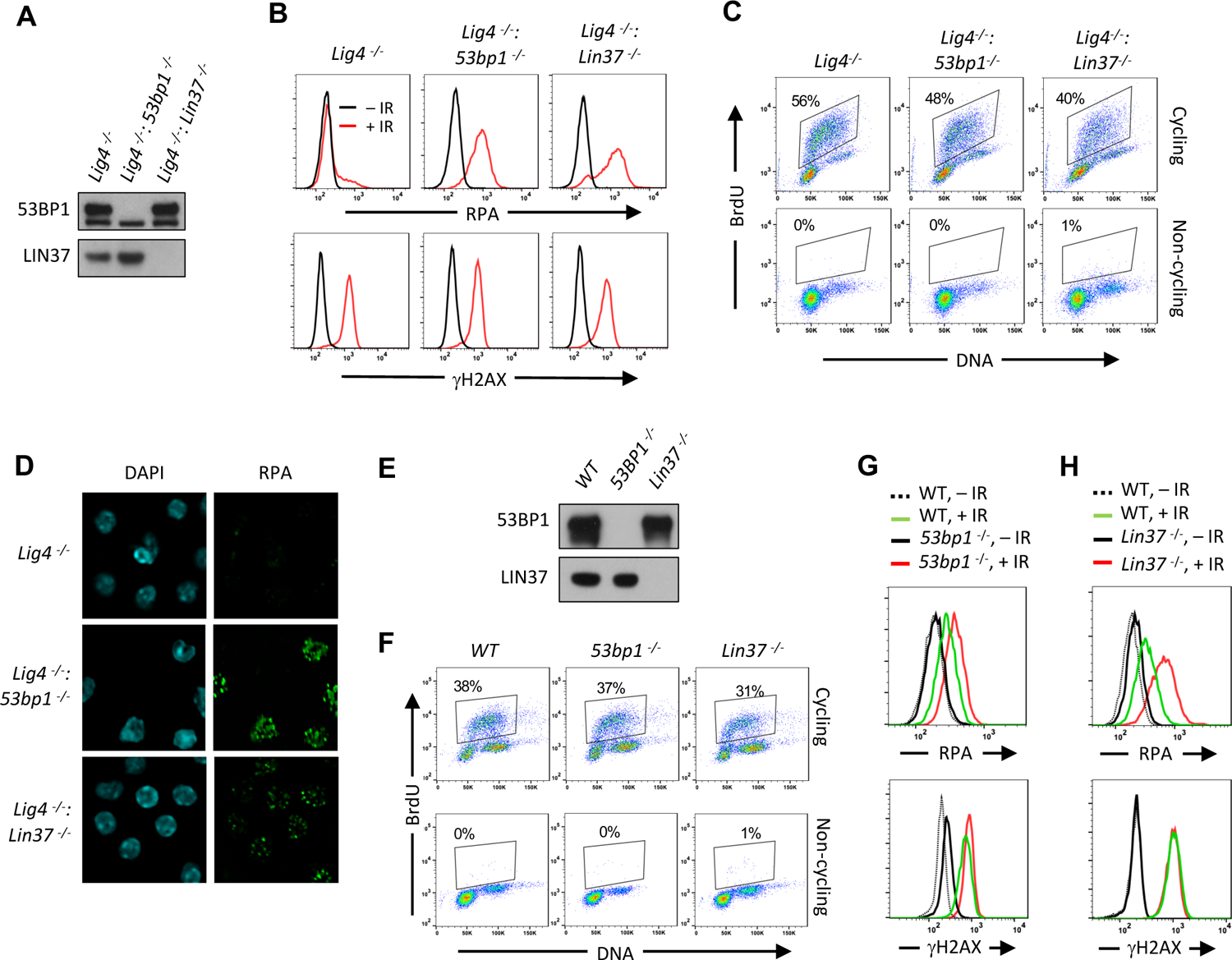
Non-cycling LIN37-deficient cells accumulate chromatin-bound RPA after IR-induced damage (A) Western blot of indicated proteins in *Lig4*^-/-^, *Lig4*^-/-^:*53bp1*^-/-^ and *Lig4*^-/-^:*Lin37*^-/-^ abl pre-B cells. (B) Flow cytometric analysis of chromatin-bound RPA (top) and γH2AX (bottom) before and after IR of non-cycling *Lig4*^-/-^, *Lig4*^-/-^:*53bp1*^-/-^ and *Lig4*^-/-^:*Lin37*^-/-^ abl pre-B cells. The experiments were repeated in two independently generated cell lines at least twice. (C) Flow cytometric analysis of cycling and non-cycling *Lig4*^-/-^, *Lig4*^-/-^:*53bp1*^-/-^ and *Lig4*^-/-^:*Lin37*^-/-^ abl pre-B cells for BrdU incorporation and DNA content (7-AAD). Percentage of cells in S-phase is indicated. (D) Representative images of IR-induced RPA foci in non-cycling *Lig4*^-/-^, *Lig4*^-/-^:*53bp1*^-/-^ and *Lig4*^-/-^:*Lin37*^-/-^ abl pre-B cells from two independent experiments. (E) Western blot analysis of indicated proteins in *WT*, *53bp1*^-/-^ and *Lin37*^-/-^ MCF10A cells. (F) Flow cytometric analysis of BrdU pulsed cycling (top) or non-cycling (bottom) *WT*, *53bp1*^-/-^ and *Lin37*^-/-^ MCF10A cells as in (C). (G, H) Flow cytometric analysis of chromatin-bound RPA (top) and γH2AX (bottom) before or after IR of non-cycling *WT* and (G) *53bp1*^-/-^ or (H) *Lin37*^-/-^ MCF10A cells.

LIN37 is a component of the DREAM complex, which along with Rb prevents cells from entering S-phase where resection of DNA ends and RPA loading normally occur at DSBs as part of HR (Weinberg 1995, Sadasivam and DeCaprio 2013, Ciccia and Elledge 2010). However, loss of LIN37 does not cause imatinib-treated abl pre-B cells to enter S-phase as evidenced by imatinib-treated *Lig4^-/-^:Lin37^-/-^* abl pre-B cells having primarily 2N DNA content and not incorporating BrdU, which would be indicative of DNA synthesis in S-phase cells (Figure 2C). If RPA is binding to chromatin at DSBs this binding should form discrete RPA IR-induced foci (IRIF) that can be visualized by immunofluorescence staining. Indeed, chromatin-bound RPA in IR treated non-cycling *Lig4^-/-^:53bp1^-/-^* and *Lig4^-/-^:Lin37^-/-^* abl pre-B cells formed discrete nuclear foci indicative of localization at DSBs (Figure 2D).

To evaluate LIN37 function in a different cell type using a distinct approach to render them non-cycling we generated LIN37- and 53BP1-deficient MCF10A human mammary epithelial cells. Upon withdrawal of epidermal growth factor (EGF) these cells stop cycling and have 2N DNA content and lack of BrdU incorporation consistent with the notion that they are in G_1_/G_0_ (Figures 2E and 2F). Non-cycling *LIN37^-/-^* and *53BP1^-/-^* MCF10A cells also exhibited increased chromatin-bound RPA after IR as compared to wild type MCF10A cells (Figures 2G and 2H), indicating that the function of LIN37 in suppressing RPA accumulation at DNA DSBs in non-cycling cells is conserved in human and mouse. As was seen above in the mouse cells, the extent of RPA accumulation in human cells lacking LIN37 after IR was greater than seen in cells lacking 53BP1, suggesting that LIN37 plays an important role in preventing DNA end resection and extensive generation of ssDNA at DSBs in non-cycling mammalian cells (Fig. 2G, 2H).

### DNA ends are resected in non-cycling LIN37-deficient cells

The CTIP nuclease is required to initiate DNA end resection during HR and DNA ends in non-cycling abl pre-B cells that are deficient in H2AX undergo resection that depends on CTIP (Sartori et al. 2007, Helmink et al. 2011). To determine whether IR-induced RPA association with chromatin in LIN37-deficient abl pre-B cells also depends on CTIP, *Lig4^-/-^:53bp1^-/-^* and *Lig4^-/-^:Lin37^-/-^* abl pre-B cells that expressed a *Ctip* gRNA were treated with doxycycline to induce Cas9 expression and maximal CTIP protein reduction, due to *Ctip* gene inactivation, prior to analysis (Figure 3A). This approach, which we refer to as “bulk gene inactivation”, allows for the depletion of proteins normally required for cell division and viability. The chromatin-bound RPA in IR treated non-cycling *Lig4^-/-^:Lin37^-/-^* and *Lig4^-/-^:53bp1^-/-^* abl pre-B cells depended on CTIP (Figure 3A), indicating that the increased level of chromatin-bound RPA after IR in cells lacking LIN37 or 53BP1 is due to DNA end resection that would lead to the generation of ssDNA.

**Figure 3.**
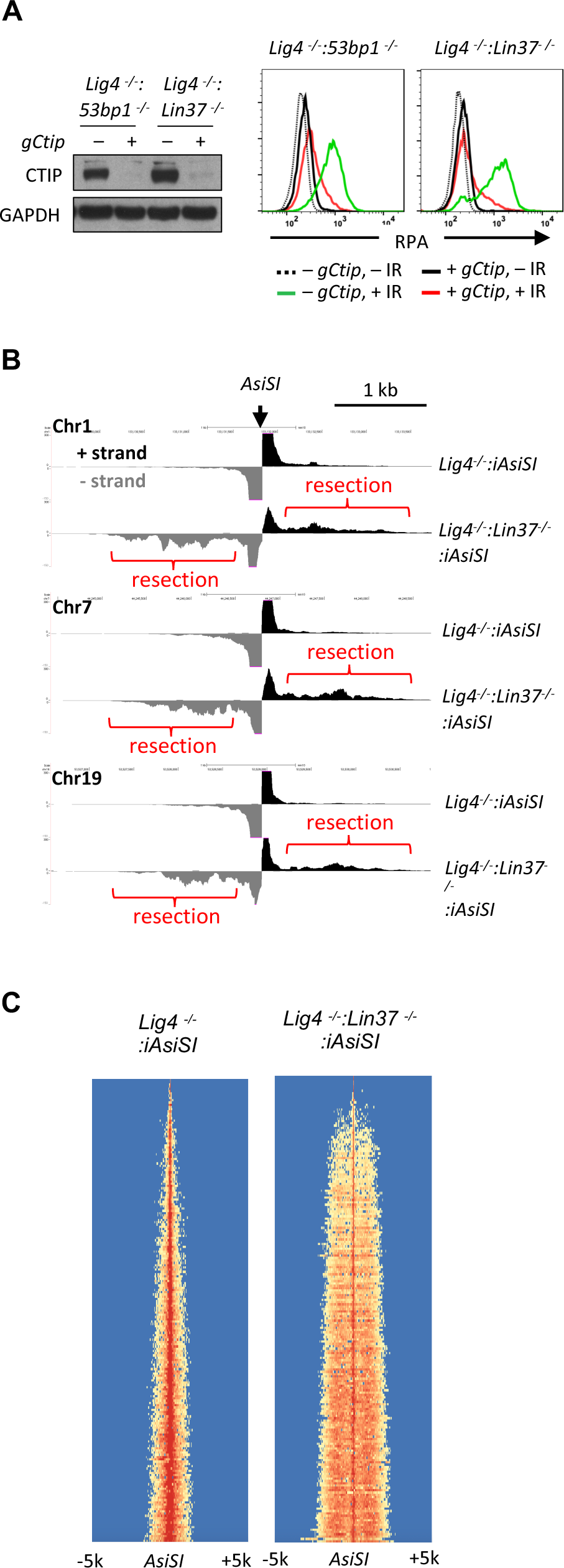
LIN37 prevents DNA end resection in non-cycling cells (A) Cas9-induced *Lig4*^-/-^:*53bp1*^-/-^ and *Lig4*^-/-^:*Lin37*^-/-^ abl pre-B cells with (+) and without (-) the *Ctip* gRNA (*gCtip*). Western blot with indicated antibodies (left) and flow cytometric analysis of chromatin-bound RPA before and after IR of the non-cycling cells (right) are shown. Representative of three experiments. (B) End-seq tracks of representative *AsiSI* sites on mouse chromosomes 1, 7 and 19 in non-cycling *Lig4*^-/-^ and *Lig4*^-/-^:*Lin37*^-/-^ abl pre-B cells. (C) The heatmaps of End-seq at *AsiSI* DSBs across the mouse genome (y axis) after *AsiSI* induction in non-cycling *Lig4*^-/-^ and *Lig4*^-/-^:*Lin37*^-/-^ abl pre-B cells. Two experiments were carried out in two independently generated *Lig4*^-/-^:*iAsiSI* and *Lig4*^-/-^:*Lin37*^-/-^:*iAsiSI* clones.

To gain direct evidence of DNA DSB end resection in LIN37-deficient abl pre-B cells, DNA End Sequencing (End-seq) was used to directly assay DNA end structures at approximately 200 *AsiSI* sites in abl pre-B cells with an inducible *AsiSI* endonuclease (*iAsiSI*) (Canela et al. 2016). End-seq allows for nucleotide resolution mapping of length and end position of DNA end resection at defined DSBs induced by a variety of endonucleases (Canela et al. 2016). The End-seq analysis revealed that that while *AsiSI* DSBs in non-cycling *Lig4^-/-^:iAsiSI* abl pre-B cells were minimally resected (<200bp), those in non-cycling *Lig4^-/-^:Lin37^-/-^:iAsiSI* abl pre-B cells were resected up to 2 kb (Figures 3B and 3C). We conclude that loss of LIN37 leads to the CTIP-dependent resection of broken DNA ends in non-cycling cells.

### LIN37 and 53BP1 are in distinct pathways of DNA end protection

53BP1 and its downstream effector proteins protect DNA ends from resection through multiple potential mechanisms (Setiaputra and Durocher 2019, Mirman and de Lange 2020, Bunting et al. 2010). To determine whether LIN37 functions in the same pathway as 53BP1, we first examined whether loss of LIN37 alters the expression levels of key proteins in the 53BP1 pathway. In this regard, Western blotting revealed that loss of LIN37 did not lead to reduction in the levels of 53BP1, RIF1 or SHLD1 proteins in cycling or non-cycling abl pre-B cells (Figures 4A and S2A). Moreover, after IR treatment, robust and near equivalent numbers of 53BP1 and RIF1 foci form in non-cycling *Lig4^-/-^* and *Lig4^-/-^:Lin37^-/-^* abl pre-B cells, demonstrating that both 53BP1 and RIF1 efficiently localize to DSBs in LIN37-deficient cells (Figures 34B, 4C, S2B and S2C).

**Figure 4.**
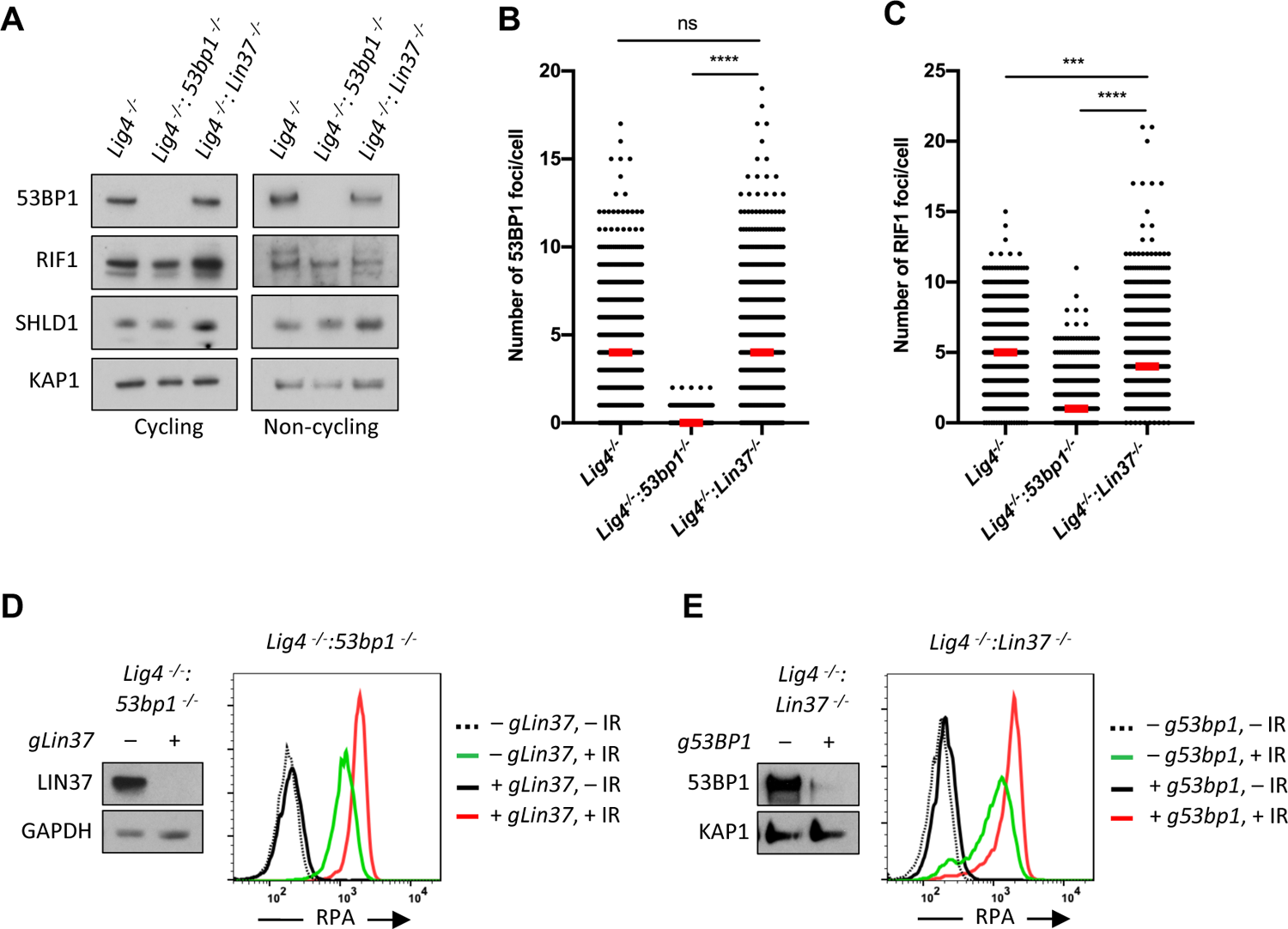
53BP1 and LIN37 have distinct DNA end protection functions (A) Western blot of indicted proteins in cycling and non-cycling *Lig4*^-/-^, *Lig4*^-/-^:*53bp1*^-/-^ and *Lig4*^-/-^:*Lin37*^-/-^ abl pre-B cells. (B, C) Quantification of 53BP1 (B) or RIF1 (C) foci after IR treatment of non-cycling *Lig4*^-/-^, *Lig4*^-/-^:*53bp1*^-/-^ and *Lig4*^-/-^:*Lin37*^-/-^ abl pre-B cells. Red bars indicate the median number of foci in each sample. More than 1000 cells were analyzed in each cell line in two independent experiments (****p<0.0001, ***p=0.0002, Mann-Whitney test). (D, E) Flow cytometric analysis of chromatin-bound RPA before and after IR of non-cycling *Lig4*^-/-^:*53bp1*^-/-^(D) or *Lig4*^-/-^:*Lin37*^-/-^ (E) abl pre-B cells after bulk gene inactivation of *Lin37* (*gLin37*) or *53bp1* (*g53bp1*), respectively. Representative of three experiments.

To test genetically whether 53BP1 and LIN37 protects DNA ends using the same or different pathways, we conducted an epistasis analysis of DNA end resection in non-cycling cells lacking LIN37, 53PB1 or both of these proteins. To this end, a *Lin37* gRNA was used to carry out bulk *Lin37* inactivation in *Lig4^-^*

*^/-^:53bp1^-/-^* abl pre-B cells (Figure 4D). Loss of LIN37 in non-cycling *Lig4^-/-^:53bp1^-/-^* abl pre-B cells led to an increase in chromatin-bound RPA after IR, as compared to non-cycling *Lig4^-/-^:53bp1^-/-^* abl pre-B cells that express LIN37 (Figure 4D). Similarly, the loss of 53BP1 in non-cycling *Lig4^-/-^:Lin37^-/-^* abl pre-B cells also led to increased chromatin-bound RPA after IR (Figure 4E). Given the additive effect of inactivation of both LIN37 and 53BP1, we conclude that in non-cycling cells 53BP1 and LIN37 are part of mechanistically distinct pathways that are both required to protect DNA ends from resection.

### LIN37 suppresses the expression of DNA end resection and HR proteins in non-cycling cells

As LIN37 participates in the formation of the DREAM transcriptional repressor we considered the possibility that Lin37 may protect DNA ends through the suppression of genes encoding pro-resection proteins (Sadasivam and DeCaprio 2013). A mutant form of LIN37, LIN37^CD^, does not associate with the DREAM complex and while DREAM lacking LIN37 binds to its target genes, it cannot repress their expression (Mages et al. 2017). To determine whether the resection of DNA ends depends on loss of DREAM transcription repressor activity, we assayed IR-induced RPA association with chromatin in non-cycling *Lig4^-/-^:Lin37^-/-^* abl pre-B cells expressing WT LIN37 or DREAM-binding defective mutant, LIN37^CD^. Expression of WT LIN37, but not LIN37^CD^, in non-cycling *Lig4^-/-^:Lin37^-/-^* abl pre-B cells decreased IR-induced chromatin-bound RPA (Figure 5A). That the extensive DNA end resection due to LIN37 deficiency cannot be rescued by expression of the LIN37 mutant that is unable to assemble into the DREAM complex suggests that LIN37 functions to protect DNA ends through its activity in the DREAM transcriptional repressor.

**Figure 5.**
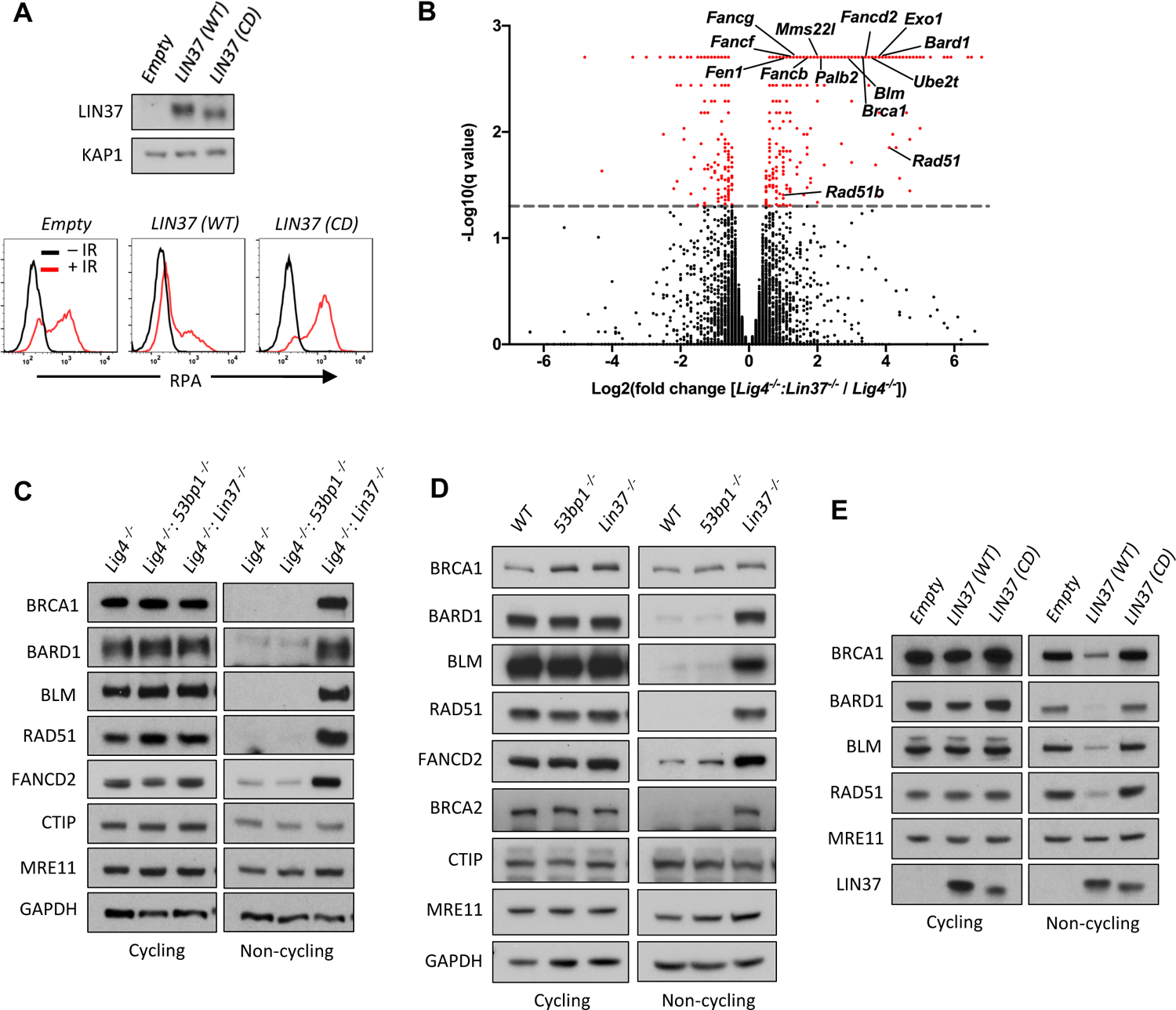
LIN37 suppresses the expression of HR protein expression in non-cycling cells (A) Western blot (top) and flow cytometric analysis for chromatin bound RPA after before or after IR (bottom) of non-cycling *Lig4*^-/-^:*Lin37*^-/-^ abl pre-B cells with empty lentivirus or lentivirus expressing wild type (WT) LIN37 or the LIN37 (CD) mutant. Representative of three experiments. (B) Volcano plot of RNA seq analysis of non-cycling *Lig4*^-/-^ and *Lig4*^-/-^:*Lin37*^-/-^ abl pre-B cells showing log2 values of the ratio of normalized transcript levels of *Lig4^-/-^*: *Lin37^-/-^* to *Lig4^-/-^* cells (X-axis) and -log10 of the q-values of fold enrichment of each gene (Y-axis). The dash line indicates q = 0.05. Genes with q values ≤ 0.05 are denoted as red dots. (C) Western blot of indicated proteins in cycling and non-cycling *Lig4*^-/-^, *Lig4*^-/-^:*53bp1*^-/-^ and *Lig4*^-/-^:*Lin37*^-/-^ abl pre-B cells. (D) Western blot analysis of indicated proteins in cycling or non-cycling *WT, 53bp1*^-/-^ and *Lin37*^-/-^ MCF10A cells. (E) Western blot of indicated proteins in cycling and non-cycling *Lig4*^-/-^:*Lin37*^-/-^ abl pre-B cells with empty lentivirus or lentivirus expressing wild type (WT) LIN37 or the LIN37 (CD) mutant.

To identify genes repressed by DREAM that may suppress end resection in non-cycling cells, we carried out RNA Seq analysis of cycling and non-cycling *Lig4^-/-^* and *Lig4^-/-^:Lin37^-/-^* abl pre-B cells. Cycling *Lig4^-/-^:Lin37^-/-^* abl pre-B cells exhibit very few (approximately 20) significant gene expression changes as compared to *Lig4^-/-^* abl pre-B cells (Table S2). In contrast, when compared to non-cycling *Lig4*^-/-^ abl pre-B cells, non-cycling *Lig4^-/-^:Lin37^-/-^* abl pre-B cells exhibited increased expression (>3-fold) of over 300 genes (Figure 5B and Table S2). We found that many genes up-regulated in non-cycling *Lig4^-/-^:Lin37^-^*

*^/-^* abl pre-B cells have functions in HR, including DNA end resection (*Brca1*, *Bard1*, *Blm*, and *Exo1*), recombination (*Brca2*, *Bard1*, *Palb2*, *Mms22l*, *Rad51*, *Rad51b*, and *Rad54b*) and DNA synthesis (*Hrob*/ *BC030867*, *Mcm3*, *Mcm4*, *Mcm5*, *Mcm7*, *Mcm8*, *Pold1*, *Pole*). A subset of Fanconi Anemia (FA) genes also exhibited increased expression in non-cycling *Lig4^-/-^:Lin37^-/-^* abl pre-B cells (Figure 5B and Table S2). These data indicate that in non-cycling abl pre-B cells LIN37-DREAM transcriptionally represses many genes that function in DNA end resection and HR.

Western blotting revealed that the levels of BRCA1, BARD1, BLM, RAD51 and FANCD2 proteins were extremely low in non-cycling *Lig4^-/-^* abl pre-B cells compared to cycling cells (Figure 5C). Consistent with their increased transcript levels, the protein levels of BRCA1, BARD1, BLM, RAD51 and FANCD2 were significantly elevated in non-cycling *Lin37^-/-^* and *Lig4^-/-^:Lin37^-/-^* abl pre-B cells as compared to WT and *Lig4^-/-^* abl pre-B cells, respectively (Figures 5C and S3). In contrast, the levels of these proteins were not altered in cycling LIN37-deficient cells (Figures 5C and S3). The increased expression of these HR proteins was also observed in non-cycling, but not cycling, *Lin37^-/-^* human MCF10A cells, indicating that LIN37 represses the expression of key DNA end resection and HR proteins to potentially limit DNA end resection in both human and murine non-cycling cells (Figure 5D).

To determine whether the ability of LIN37 to repress expression of these key HR proteins in non-cycling cells required LIN37 to function in the context of the DREAM complex, we again used the LIN37^CD^ mutant which cannot form a transcriptionally repressive DREAM complex. We found that expression of wild type LIN37, but not LIN37^CD^ in non-cycling *Lig4^-/-^:Lin37^-/-^* abl pre-B cells, led to a decrease in the levels of BRCA1, BARD1, BLM, RAD51 and FANCD2 proteins (Figure 5E). Of note is that the expression of the MRE11 and CTIP nucleases were not affected by loss of LIN37 in cycling or non-cycling cells (Figures 5C and S3). We conclude that in non-cycling cells, LIN37, as part of the DREAM transcriptional repressor complex, represses genes encoding many proteins that function in DNA end resection and DSB repair by HR.

### DNA resection and HR machinery are functional in non-cycling LIN37-deficienct cells

We performed another whole genome CRISPR/Cas9 reverse genetic screen to identify genes that mediate the extensive DNA end resection that occurs in non-cycling LIN37-deficient abl pre-B cells. To this end we identified gRNAs enriched in IR treated non-cycling *Lig4^-/-^:Lin37^-/-^* abl pre-B cells that had low levels of chromatin-bound RPA, indicating reduced DNA end resection despite having no LIN37 (Table S3). As expected, gRNAs to *Rbbp8* (encodes CTIP) and to *Mre11b* (encodes MRE11) were enriched in these RPA low cells, in agreement with their nucleolytic roles in resection and emphasizing the validity of our screen (Table S3). In addition, we isolated gRNAs to many genes encoding DNA end resection and HR proteins that are normally repressed by LIN37, including *Brca1*, *Bard1*, *Blm* and several FA genes, indicating that the expression of these proteins brought about by LIN37-deficiency in non-cycling cells promotes the extensive DNA end resection that is observed in these cells (Table S3). Indeed, bulk inactivation of BRCA1, BARD1, BLM or FANCD2 significantly reduced the level of chromatin-bound RPA after IR in non-cycling *Lig4^-/-^:Lin37^-/-^* abl pre-B cells (Figure 6A). In striking contrast, the loss of these proteins had no effect on RPA localization to DSBs after IR treatment of non-cycling *Lig4^-/-^:53bp1^-/-^* abl pre-B cells, further emphasizing that LIN37 and 53BP1 are part of mechanistically distinct pathways that prevent DNA end resection and ssDNA generation in non-cycling cells (Figures 6A).

**Figure 6.**
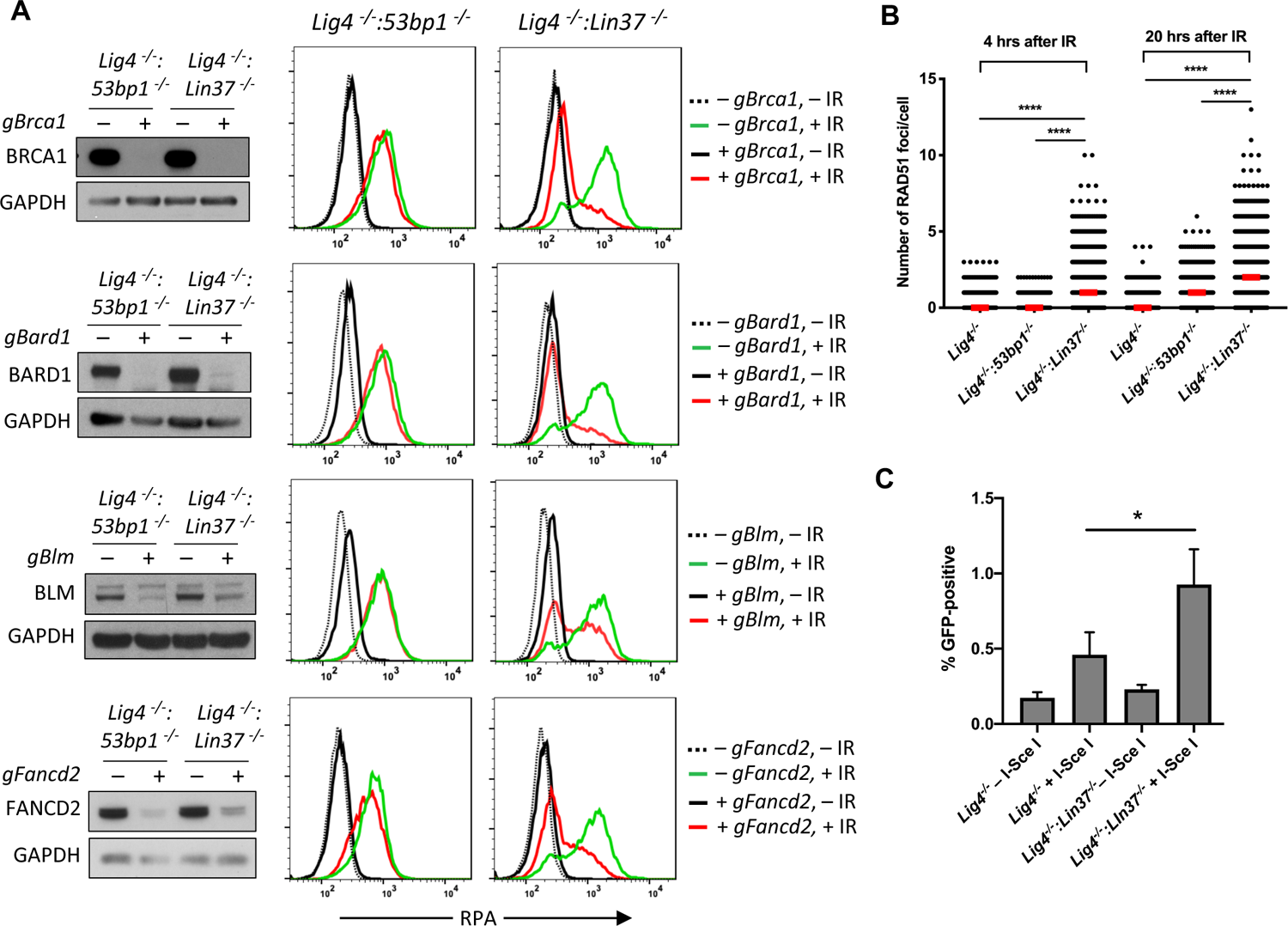
LIN37 Prevents resection and HR through suppressing HR protein expression in non-cycling cells (A) Flow cytometric analysis of chromatin-bound RPA before and after IR of non-cycling *Lig4*^-/-^:*53bp1*^-/-^ or *Lig4*^-/-^:*Lin37*^-/-^ abl pre-B cells with or without indicated gRNAs following Cas9 induction for bulk gene inactivation. Representative experiments of three. (B) Quantification of RAD51 foci in non-cycling *Lig4*^-/-^, *Lig4*^-/-^:*53bp1*^-/-^ and *Lig4*^-/-^:*Lin37*^-/-^ abl pre-B cells 4 and 20 hours after IR. Red bars indicate the median number of RAD51 foci in each sample of more than 1000 cells analyzed for each cell line. Representative of two independent experiments (****p<0.0001, Mann-Whitney test). (C) Flow cytometric analysis of HR-mediated DSB repair in non-cycling *Lig4*^-/-^ and *Lig4*^-/-^:*Lin37*^-/-^ abl pre-B cells using the HPRT-DR-GFP reporter. The percentage of GFP-positive cells is shown. Error bars are ± SEM from 3 experiments (*p=0.0124, t-test).

LIN37-deficient non-cycling abl pre-B cells also exhibited induction of *Brca2*, *Palb2* and *Rad51,* which function to replace RPA with RAD51 on ssDNA to form RAD51 nucleofilaments at DSBs during HR (Figures 5C, 5D and S3). Indeed, there was a significant increase in the number of RAD51 foci in non-cycling *Lig4^-/-^:Lin37^-/-^* abl-pre-B cells after IR as compared to *Lig4^-/-^* abl-pre-B cells (Figures 6B and S4). Once formed, a RAD51 nucleofilament will initiate a homology search and HR-mediated DSB repair (Prakash et al. 2015). Indeed, using the HPRT-DR-GFP reporter for DSB repair by HR, we observed an increase in homology-mediated DSB repair in non-cycling *Lig4^-/-^:Lin37^-/-^* abl-pre-B cells as compared to *Lig4^-/-^* abl-pre-B cells (Figure 6C) (Pierce et al. 2001). We conclude that in the absence of LIN37, the extensive DNA end resection can lead to HR-mediated DSB repair in non-cycling cells. Moreover, this occurs in the presence of 53BP1.

### LIN37 uniquely prevents DNA end resection in quiescent G_0_ cells

We find that LIN37 is required to prevent DNA end resection and subsequent HR steps in non-cycling cells that have 2N DNA content and do not incorporate BrdU indicating that they are in G_0_ or G_1_. To distinguish between G_0_ and G_1_ we assayed non-cycling imatinib-treated abl pre-B cells and EGF-deprived MCF10A cells for expression of the cyclin dependent kinase, CDK4, and phospho-CDK4, which along with CDK6 is required for non-cycling quiescent G_0_ cells to move into G_1_ (Figures 7A and 7B) (Malumbres and Barbacid 2001, Pennycook and Barr 2020). Both non-cycling abl pre-B cells and MCF10A cells had low levels of CDK4 and phsopho-CDK4 indicative of them being in G_0_ (Figures 7A and 7B). Rb suppresses genes encoding proteins required for G_0_ cells to transit to G_1_ and then into S-phase (Weinberg 1995, Pennycook and Barr 2020, Sadasivam and DeCaprio 2013). The phosphorylation of Rb by CDK4 or CDK6 leads to its inactivation and the transit of cells from G_0_ to G_1_ (Weinberg 1995, Pennycook and Barr 2020). Phospho-Rb was not detected by western blotting of imatinib-treated abl pre-B cells or MCF10A cells deprived of EGF indicating that these cells are in G_0_ (Figures 7A and 7B). PCNA, which is expressed early in the transition from G_1_ to S was also not detected in these cells (Figures 7A and 7B). Together these data indicate that imatinib treated abl pre-B cells and MCF10A cells deprived of EGF have exited the cell cycle and entered G_0_, also known as quiescence, and LIN37 is required to protect DNA ends from resection in these cells.

**Figure 7.**
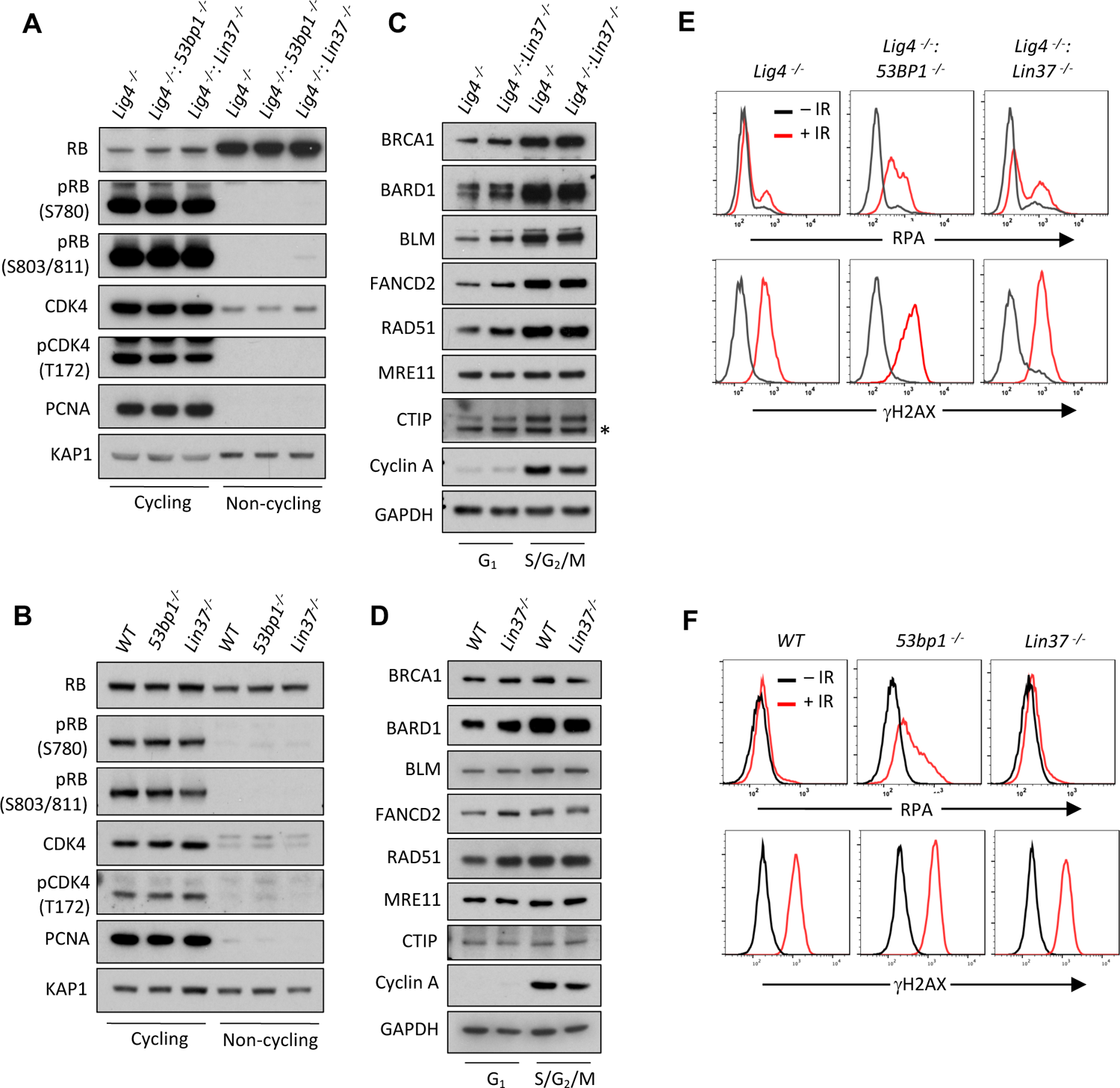
LIN37 function in DNA end protection is restricted to G_0_ (A, B) Western blot of indicated proteins in cycling and non-cycling abl pre-B cells (A) or MCF10A cells (B). (C, D) Western blot of indicated proteins in cycling G_1_ or S/G_2_/M abl pre-B cells (C) or MCF10A cells (D), isolated by flow cytometric cell sorting based on the PIP-FUCCI reporter. Representative of two independent experiments. Asterisk indicates non-specific recognizing bands. (E, F) Flow cytometric analysis of chromatin bound RPA and γH2AX before and after IR treatment of G_1_ phase *Lig4^-/-^*, *Lig4^-/-^:53bp1*^-/-^ and *Lig4^-/-^:Lin37*^-/-^ abl pre-B cells (E) or *WT*, *53bp1*^-/-^ and *Lin37*^-/-^ MCF10A cells (F). Representative of three experiments.

Cycling cells in G_1_-phase also rely on NHEJ to repair DNA DSBs and we next asked whether LIN37 also functions to protect DNA ends from resection in these cells. To this end we employed the PIP-FUCCI cell cycle sensor to isolate G_1_ and S/G_2_/M populations from cycling *Lig4^-/-^*, *Lig4^-/-^:53bp1^-/-^* and *Lig4^-/-^:Lin37^-/-^* abl pre-B cells and wild type, *53bp1^-/-^* and *Lin37^-/-^* MCF10A cells by flow cytometric cell sorting (Grant et al. 2018). The effectiveness of this purification was evidenced by the absence of Cyclin A in G_1_-phase cells (Figures 7C and 7D) (Henglein et al. 1994). Western blot analysis revealed that in contrast to G_0_ cells, G_1_ cells expressed detectable levels of the HR proteins BRCA1, BARD1, BLM, FANCD2 and RAD51. Moreover, loss of LIN37 did not lead to a significant increase in the levels of these proteins in G_1_-phase cells isolated from proliferating populations (Figures 7C and 7D).

We next asked whether LIN37 functions to protect DNA ends from extensive end resection in cycling G_1_-phase cells. To do this, we incubated proliferating *Lig4^-/-^*, *Lig4^-/-^:53bp1^-/-^* and *Lig4^-/-^:Lin37^-/-^* abl pre-B cells and wild type, *53bp1^-/-^* and *Lin37^-/-^* MCF10A cells in media containing EdU, which was incorporated into newly-synthesized DNA allowing for the detection of S-phase cells. After IR, RPA association with DNA DSBs was assayed by flow cytometry in G_1_-phase cells, which were identified as cells being EdU-negative and having 2N DNA content (Figures S5A and S5B). In contrast to G_0_ cells, which are dependent on 53BP1 and Lin37 for DNA end protection, G_1_ cells appear to depend primarily on 53BP1 as evidenced by the substantial increase in RPA association with chromatin in IR treated 53BP1-deficient G_1_-phase cells as compared to LIN37-deficient G_1_-phase cells (Figures 7E and 7F). We conclude that LIN37 functions to prevent DNA end resection primarily in quiescent G_0_ cells while 53BP1 functions in both proliferating G_1_ and quiescent G_0_ phase cells.

## Discussion

Rb and the DREAM complex are transcriptional repressors that silence the expression of genes required to promote cell cycle, driving cells to exit the cell cycle an enter G_0_ or quiescence, where most cells in the human body reside (Mages et al. 2017, Weinberg 1995). Quiescent G_0_ cells and cycling cells in G_1_-phase have 2N DNA content and rely on NHEJ to repair DNA DSBs and maintain genome stability. 53BP1 and its downstream effectors are required to promote NHEJ in G_0_ and G_1_ cells by antagonizing DNA end resection and ssDNA generation (Mirman and de Lange 2020, Setiaputra and Durocher 2019). However, here we show that in G_0_ cells, DNA end protection also requires LIN37, a component of the DREAM complex. Moreover, LIN37 function also prevents aberrant homology-mediated joining in G_0_ cells.

LIN37 association with DREAM is required for its transcriptional repressor function, but is not required for the binding of the complex to its target genes (Mages et al. 2017). In addition to genes encoding proteins that promote cell cycle, DREAM complex binding sequences, termed cell cycle genes homology regions (CHRs), are also found in the promoter regions of many HR genes (Mages et al. 2017, Muller et al. 2014). Several lines of evidence presented here demonstrate that in G_0_ cells LIN37 functions through DREAM transcriptional repression to protect DNA ends from resection. Loss of LIN37 in G_0_ cells leads to the expression of many HR genes that promote DNA end resection such as *Brca1*, *Bard1*, *Blm* and *Fancd2* (Figure 5B and Table S2). Moreover, this increased gene expression leads to a significant increase in the levels of these proteins in G_0_ cells (Figures 5C, 5D and S3). Loss of BRCA1, BARD1, BLM or FANCD2 in LIN37-deficient G_0_ cells prevents DNA end resection demonstrating that each of these function of each of these proteins is required to promote DNA end resection in these cells (Figure 6A). Finally, while the expression of wild type LIN37 in LIN37-deficient G_0_ cells prevents the expression of HR genes and DNA end resection the expression of LIN37^CD^, which cannot participate in forming a functional DREAM repressor complex, does not (Figure 5E).

LIN37-DREAM and 53BP1 function in distinct ways to protect DNA ends from aberrant resection. This notion is supported by the additive effect on resection in *Lig4^-/-^* G_0_ abl pre-B cells lacking both 53BP1 and LIN37, as compared to those lacking 53BP1 or LIN37 in (Figures 4D and 4E). Resection in G_0_ LIN37-deficienct cells relies on BRCA1, BARD1, BLM and FANCD2 as evidenced by the lack of DNA end resection in LIN37-deficienct cells that have lost expression of any of these proteins through bulk gene inactivation (Figure 6A). In contrast, 53BP1-deficienct G_0_ cells express low levels of BRCA1, BARD1, BLM and FANCD2 and bulk inactivation of the genes encoding these proteins does not impact DNA end resection in these cells (Figure 6A). Thus, in G_0_ cells LIN37 prevents BRCA1-dependent DNA end resection whereas 53BP1 protects DNA ends from BRCA1-independent resection pathways.

Cycling cells in G_1_-phase also must repair DSBs by NHEJ and thus need to protect DNA ends from excess resection. In contrast to what we observe in G_0_ cells, the loss of LIN37 in cycling cells in G_1_-phase does not lead to alterations in HR protein levels or aberrant resection of DNA ends as evidenced by increased RPA association at DSBs (Figure 7). In G_1_-phase cells, 53BP1-RIF1 prevents BRCA1-CTIP from associating with DSBs and promoting resection (Escribano-Diaz et al. 2013, Chapman et al. 2013). In contrast, in S/G_2_-phase cells this regulatory balance is tipped with BRCA1 inhibiting RIF1 association with DSBs, which would otherwise antagonize resection (Escribano-Diaz et al. 2013, Chapman et al. 2013). Like G_1_-phase cells, in G_0_ cells 53BP1 functions to protect DNA ends from resection. However, upon the loss of LIN37 in G_0_ cells the expression of HR proteins, including BRCA1, does not inactivate the 53BP1 pathway. Indeed, we find robust 53BP1-RIF1 association with DSBs in LIN37-deficient G_0_ cells and the loss of 53BP1 in these cells leads to increased RPA association at DSBs indicating that 53BP1 functions in DNA end protection in these cells (Figures 4B-E, S2B and S2C). We speculate that the genetic program activated by the loss of LIN37 in G_0_ cells leads to the activation of pathways that regulate the balance of anti-resection 53BP1 activities and pro-resection BRCA1 activities in a manner that favors DNA end resection. Although the identity of these pathways and the manner in which they function is unknown, they do not prevent 53BP1-RIF1 association at DSBs in G_0_ cells.

While LIN37-DREAM and 53BP1 both inhibit DNA end resection in G_0_ cells, LIN37-DREAM has additional activities in promoting genome stability by preventing resected DNA ends from progressing in HR-mediated DSB repair. LIN37-deficient G_0_ cells express BRCA2, PALB2 and RAD51 that convert RPA-coated ssDNA at broken DNA ends to RAD51 nucleofilaments that can mediate homology mediated repair (Figures 5B-D, 6B, 6C, S3, S4 and Table S2), which in G_0_ cells could lead to aberrant homology-mediated DNA end joining and chromosomal aberrations such as deletions and translocations. Thus, in quiescent cells in the body, LIN37-DREAM promotes genome stability by both antagonizing DNA end resection and preventing the aberrantly homology-mediated joining of resected DNA ends.

## Supporting information

RNA seq in Lig4 and Lig4_Lin37 cells

RPA screen in Lig4 KO cells

RPA screen in LIg4-Lin37 KO cells

## Acknowledgement

B.P.S. is supported by National Institutes of Health grants R01 AI047829 and R01 AI074953. J.K.T. is supported by National Institutes of Health grants R01 CA095641 and R01 GM064475.

## Author Contributions

B.C., A.N, J.K.T and B.P.S. designed the study. And B.C. conducted all the experiments, except the ones specified below. Y.W. assisted genome-scale guide RNA CRISPR/Cas9 screen and performed data analysis. D.Z and F.F. performed imaging experiments. A.T.T., N.Z. and W.W. conducted End Seq and performed data analysis. A.B., C.C. and W.F. performed additional experiments. B.C., J.K.T and B.P.S wrote the manuscript with the help of A.N. All authors reviewed the manuscript.

## Materials and Methods

### Cell lines and cell culture

Abelson virus-transformed pre-B cells (abl pre-B cells) were generated as described previously(Bredemeyer et al. 2006). DNA Ligase 4-deficient (*Lig4^-/-^*) abl pre-B cells were generated by deleting the LoxP site-flanked *Lig4* coding sequence by expressing Cre recombinase(Helmink et al. 2011). WT and *Lig4^-/-^* abl pre-B cells and MCF10A human mammary epithelial cells used in this study all contain pCW-Cas9 (Addgene# 50661), which has a FLAG-tagged *Cas9* cDNA under the control of a doxycycline-inducible promoter. Intracellular staining with anti-FLAG and flow cytometry were used to identify clones homogeneously expressing FLAG-CAS9 after doxycycline treatment.

*53bp1*^-/-^, *Lin37*^-/-^, *Lig4^-/-^*:*53bp1*^-/-^ and *Lig4^-/-^*:*53bp1*^-/-^ cell lines were made by transiently transfecting *53bp1* or *Lin37* guide RNAs (gRNAs) in the pX330 vector (Addgene# 42230) into *WT* or *Lig4^-/-^* cells followed by subcloning by limited dilution. Knockout clones were verified by western blotting for loss of expression of the proteins encoded by genes targeted by the gRNAs. Abl pre-B cells were cultured in Dulbecco’s Modified Eagle Medium (DMEM) supplemented with 10% fetal bovine serum (FBS) and 0.4% beta-mercaptoethanol, 100 U/ml penicillin/streptomycin, 1mM sodium pyruvate, 2mM L-glutamine,1X nonessential amino acids. MCF10A cells were cultured in DMEM/F12 supplemented with 5% horse serum, 20ng/ml EGF, 0.5μg/ml hydrocortisone, 100ng/ml cholera toxin, 10μg/ml insulin and 100U/ml penicillin-streptomycin.

To render cells non-cycling, abl pre-B cells were treated with 3 μM imatinib (Selleck Chemicals, S2475) for 2 (for chromatin-bound RPA assay) or 3 (for gene expression analysis) days and MCF10A cells were grown in EGF-free media for 2 days. At least two independent knockout clones of each gene in each cell type were generated and used in the experiments in this study. Knockout clones were verified by western blot for lack of expression of the targeted proteins.

### Chromatin-bound RPA assay

The chromatin-bound RPA flow cytometry assay was carried out as described previously with modifications(Forment, Walker, and Jackson 2012). Briefly, cells were washed with FACS wash (2% FBS in 1X PBS) followed by pre-extraction in Triton X-100 on ice for 10 minutes (0.05% for imatinib-treated abl pre-B cells, 0.2% for proliferating abl pre-B cells and 0.5% for MCF10A cells). Cells were washed again with FACS wash and fixed in BD Cytofix/Cytoperm at room temperature for 10 minutes. After fixation, cells were incubated with anti-RPA32 antibody (Cell Signaling Technology, 2208S, 1:500) and anti-phospho-H2AX (S139) (Millipore Sigma, 05-636, 1:1000) in 1X BD Perm/Wash buffer at room temperature for 2 hours, followed by staining with Alexa Fluor 488 Goat anti-rat IgG (BioLegend, 405418, 1:500) and Alexa Fluro 647 Goat anti-moue IgG (BioLegend, 405322, 1:500) at room temperature for 1 hour. 20 μl of 7-AAD (BD Pharmingen, 559925) was added to each sample before resuspending cells in 300 μl of 1X PBS. RPA32 and phospho-H2AX(S139) levels were analyzed on a BD LSRFortessa Flow Cytometer.

For analysis of G_1_ cells in a proliferating population, cells were pulsed with 10 μM EdU for 1 hour prior to irradiation. Irradiated cells were kept in EdU-containing media during the course of the experiments and processed as described above. Following RPA32 and phosphor-H2AX(S139), click-IT chemistry was performed as per manufacturer’s instructions.

Non-cycling abl pre-B cells were exposed to 15 Gy of IR and analyzed 18 hours after irradiation. Cycling abl pre-B cells were analyzed 3 hours after 5 Gy of IR. Non-cycling MCF10A were exposed to 30 Gy of IR and analyzed 4 hours after irradiation. Cycling MCF10A cells were analyzed 6 hours after 25 Gy IR.

### Genome-wide guide RNA CRISPR/Cas9 screen

*Lig4^-/-^* or *Lig4^-/-^:Lin37^-/-^* abl pre-B cells were transduced with lentiviral mouse genome-wide CRISPR gRNA library V2 (Addgene #67988) by centrifuging a cell and viral supernatant mixture (supplemented with 5μg/ml polybrene) at 1,800 rpm for 90 minutes. BFP-positive (stably transduced) cells were isolated on BD FACSAria II Cell Sorter, treated with 3μg/ml doxycycline for 7 days followed by treatment with 3μM imatinib for 2 days. 18 hours after exposing to 20 Gy IR, cells were processed as described above for the chromatin-bound RPA flow cytometry assay and analyzed on a BD FACSAria II Cell Sorter. Cells with high (top 10%), low (bottom 10%) RPA staining, as well as unsorted cells were collected, and genomic DNA of these cells were harvested for amplification of gRNAs.

To generate an Illumina sequencing library, gRNAs in the selected cells were first amplified using primers pKLV lib330F and pKLV lib490R and the program “98°C/5 minutes - [98°C/15 seconds - 60°C/15 seconds - 68°C/1 minute]x18 - 68°C/5 minutes”. The resulting PCR products were used as the template for additional PCR amplification using primers PE.P5_pKLV lib195 Fwd and P7 index180 Rev and the program “94°C/5 minutes - [94°C/15 seconds - 60°C/30 seconds - 68°C/20 seconds]x10 - 68°C/5 minutes” to add Illumina HiSeq adapters and indexes (RPI5:CACTGT, RPI6: ATTGGC. RPI12: TACAAG). The final PCR products (∼300 bp) were resolved in 1.5% agarose gel and purified by QIAQuick Gel Purification Kit (QIAGEN). Purified DNA was sequenced on Illumina HiSeq2500 system to determine gRNA representation in each sample (50bp single-end reads).

Raw fastq files were demultiplexed by the Genomics and Epigenomics Core Facility of the Weill Cornell Medicine Core Laboratories Center. The gRNA sequence region was then retrieved from the sequencing data using Seqkit(Shen et al. 2016) and mapped to the gRNA sequence library(Koike-Yusa et al. 2014, Tzelepis et al. 2016). The number of reads of each library sequence was counted and then normalized as follows(Shalem et al. 2014):

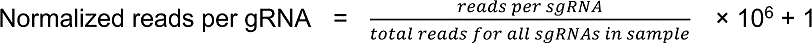

Lastly, the enrichment score of a gRNA was calculated as a ratio of normalized reads of the gRNA in two samples.

pKLV lib330F: AATGGACTATCATATGCTTACCGT pKLV lib490R CCTACCGGTGGATGTGGAATG

PE.P5_pKLV lib195 Fwd: AATGATACGGCGACCACCGAGATCTGGCTTTATATATCTTGTGGAAAGGAC

P7 index180 Rev: CAAGCAGAAGACGGCATACGAGAT*<u>INDEX</u>*GTGACTGGAGTTCAGACGTGTGCTCTTCCGATCCAGACTGCCTTGGGAAAAGC

### Bulk gene inactivation

To inactivate *53bp1*, *Lin37*, *CtIP*, *Brca1*, *Bard1*, *Blm* or *Fancd2* in bulk cells populations, cells were transduced with lentivirus pKLV-gRNAs by mixing cell suspensions with viral supernatant supplemented with 5μg/ml polybrene and 3μg/ml doxycline and spinning at 1,800 rpm for 90 minutes. Stably transduced cells were sorted 3 days after transduction and sorted cells were kept in growth media with 3μg/ml doxycline for additional 24 hours before being subjected to analysis.

### Plasmid constructs

pCW-Cas9 was a gift from Eric Lander & David Sabatini (Addgene plasmid # 50661)(Wang et al. 2014). pX330-U6-Chimeric_BB-CBh-hSpCas9 was a gift from Feng Zhang (Addgene plasmid # 42230)(Cong et al. 2013). pKLV-U6gRNA(BbsI)-PGKpuro2ABFP was a gift from Kosuke Yusa (Addgene plasmid # 50946)(Koike-Yusa et al. 2014). pLenti-CMV-Blast-PIP-FUCCI was a gift from Jean Cook (Addgene plasmid # 138715) (Grant et al. 2018).

For LIN37 reconstitution experiments, Lin37 WT or CD mutant were cloned into tetracycline-inducible TRE-Thy1.1 lentriviral vector to generate TRE-Lin37 (WT)-Thy1.1 or TRE-Lin37 (CD)-Thy1.1. WT Lin37 was amplified from Lin37 cDNA BC013546 (transOMIC) using primers Lin37 iso1_5’XhoI_S and Lin37 3’NotI_AS. To generate Lin37 CD mutant, Lin37 iso1_5’XhoI_S and Lin37 CD1_AS, and Lin37 CD2_S and 3’NotI_AS were first used to generate two cDNA fragments containing mutations in CD1 and CD2. The resulting PCR products were used in an overlapping PCR that annealed the two fragments to generate the full length Lin37 (CD) mutant.

Lin37 iso1_5’XhoI_S: GCCCTCGAGATGTTCCCGGTAAAGGTGAAAGTGG Lin37 3’NotI_AS: GCCGCGGCCGCTCACTGCCGGTCATACATCTCCCGT Lin37 CD1_AS: TACAGTGGTGTGTTCTCACTGAACTGGGCCAAGTCCACAGCCCCG GCAAATAGCTTGATC

Lin37 CD2_S: ACTTGGCCCAGTTCAGTGAGAACACACCACTGTACCCCATCGCCGG CGCCTGGATGCGCA

### End-Seq

End-seq was performed as previously described, using 20×10^6^ abl pre-B cells harboring TET-inducible AsiSI-ER fusion treated with 3μM imatinib(Canela et al. 2016). Briefly, cells were embedded in agarose plugs, lysed, and treated with proteinase K and RNase A. The agarose-embedded genomic DNA was then blunted using ExoVII (NEB) and ExoT (NEB). Blunted DNA ends were A-tailed using Klenow exo-(NEB), and a biotinylated hairpin adaptor BU1 was ligated. After adaptor ligation, DNA was recovered after plug melting and treatment with beta-agarase. DNA was sheared to a length between 150 and 200 bp by sonication (Covaris), and biotinylated DNA fragments were purified using streptavidin beads (MyOne C1, Invitrogen). Following streptavidin capture, the newly generated ends were end repaired using T4 DNA polymerase, Klenow fragment, and T4 polynucleotide kinase; A-tailed with Klenow exo-fragment (15 U); and finally ligated to hairpin adaptor BU2 using the NEB Quick ligation kit. After the second adaptor ligation, libraries were prepared by first digesting the hairpins on both adapters with USER enzyme (NEB) then PCR amplified for 16 cycles using TruSeq index adapters. All libraries were quantified using qPCR. Sequencing was performed on the Illumina NextSeq500 (75 bp single-end reads).

END-seq reads were aligned to the mouse genome (GRCm38p2/mm10) using Bowtie v1.1.2 with parameters (-n 3 -k 1 -l 50) and alignment files were generated and sorted using SAMtools and BEDtools(Li et al. 2009, Quinlan and Hall 2010, Langmead et al. 2009). Heatmap was plotted using heatmap.2 of gplots package in R.

BU1: 5’-Phos-GATCGGAAGAGCGTCGTGTAGGGAAAGAGTGUU[Biotin-dT]U[Biotin-dT]UUACACTCTTTCCCTACACGACGCTCTTCCGATC*T-3’ [*phosphorothioate bond] BU2: 5’-Phos-GATCGGAAGAGCACACGTCUUUUUUUUAGACGTGTGCTCTTCCGA TC*T-3’ [*phosphorothioate bond]

### Antibodies for western blotting

The following antibodies were used for western blot analysis: 53BP1 (Bethyl Laboratories, A300-272A, 1:3000), LIN37 (Santa Cruz Biotechnology, sc-515686, 1:200), BLM (Bethyl Laboratories, A300-572A, 1:2000), BRCA1 for mouse (R&D Systems, gift from Dr. Andre Nussenzweig, NCI, 1:1000)(Zong et al. 2019), BRCA1 for human (Millipore Sigma, 07-434, 1:1000), RAD51(Millipore Sigma, ABE257, 1:2000), BARD1 (Thermo Fisher Scientific, PA5-85707, 1:1000), CtIP (gift from Dr. Richard Baer, Columbia University, New York), 1:1000), MRE11 (Novus Biologicals, NB100-142), RIF1 (Abcam, ab13422, 1:500), SHLD1/C20orf196 (Thermo Fisher Scientific, PA5-559280, 1:200), GAPDH (Sigma, G8795, 1:10000), KAP1 (Genetex, GTX102226, 1:2000), FANCD2 (R&D Systems, MAB93691, 1:1000), BRCA2 for human (Proteintech, 19791-1-AP, 1:500), Rb1 (Thermo Fisher Scientific, LF-MA0173, 1:1000), Phospho-Rb (Ser780) (Cell Signaling Technology, 8180T, 1:1000), Phospho-Rb (Ser807/811) (Cell Signaling Technology, 8516T, 1:1000), PCNA (Bethyl Laboratories, A300-276A, 1:3000).CDK4: (Novus Biologicals, NBP1-31308, 1:1000), CDK4 (phosphor Thr 172): (GeneTex, GTX00778, 1:1000).

### RNA sequencing analysis

RNA was purified from cycling *Lig4^-/-^* or *Lig4^-/-^:Lin37^-/-^* cells treated with imatinib for 72 hours (two biological replicates each) using RNeasy mini kit (Qiagen). RNAseq libraries were prepared and directional RNA sequencing of 2 × 50 bp performed at the Transcriptional Regulation & Expression Facility at Cornell University using NextSeq 500 sequencer. The raw fastq reads were first processed with Trim-Galore (Barbraham Institute). The filtered reads were then aligned to GRCm38 reference genome with ENSEMBL annotations using Spliced Transcripts Alignment 2.7 (STAR 2.7)(Dobin et al. 2013). Differential expression was computed using DESeq2 (Bioconductor) and a FDR 0.05 cutoff was used to identify sets of differentially expressed genes (Love, Huber, and Anders 2014).

### Ionizing radiation-induced foci formation (IRIF) assay

*Lig4^-/-^, Lig4^-/-^:Lin37^-/-^* and *Lig4^-/-^:53bp1^-/-^* abl pre-B cells were treated or not with 3 μM imatinib for 48 hours. Thereafter, cells were pulsed with 10 μM EdU (Invitrogen) for 30 minutes and subjected to 10 Gy irradiation. For the detection of RAD51 foci, irradiated cells were allowed to recover for 4 or 20 hours, at which point they were immobilized on slides pre-coated with CellTak (Corning) and briefly pre-extracted (20 mM HEPES, 50 mM NaCl, 3 mM MgCl_2_, 0.3 M sucrose, 0.2% Triton X-100) on ice for 15 seconds to remove soluble nuclear proteins. Extracted samples were then fixed (4% paraformaldehyde), permeabilized (0.5% Triton X-100 in PBS), incubated with anti-RAD51 primary antibody (Abcam, ab176458, 1:250). Alternatively, irradiated (10 Gy) cells were allowed to recover for 1 hour prior to fixation without a preceding pre-extraction step, and subsequently incubated with primary antibodies recognizing 53BP1 (Novus Biologicals, NB100-305,1:1000) or RIF1 (gift from Davide Robbiani (Rockefeller University, New York),1:5000). In all cases, IRIFs were visualized by incubating samples with Alexa Fluor 555-conjugated secondary antibodies (Invitrogen). Where indicated, click-IT chemistry was performed as per manufacturer’s instructions. Finally, DNA was counterstained with DAPI (Thermo Fisher Scientific). Immunofluorescence images were captured at 40X magnification on a Lionheart LX automated microscope (BioTek Instruments, Inc.). Quantification of IRIF was performed using the Gen5 spot analysis software (BioTek).

### gRNAs

Bulk gene inactivation or stable knockout mutants was achieved by CRSPR/Cas9 using the following gRNAs: *g53bp1* (GAACCTGTCAGACCCG ATC); *gLin37* (AAGCTATTTGACCGGAGTG); *gCtip* (ATTAACCGGCTACGA AAGA); *gBrca1* (GTCTACATTGAACTAGGTA); *gBard1* (AAATCGTAAAGGCT GCCAC); *gBlm* (GATTTAACGAAGGAATCGG); *gFancd2* (TCTTGTGATGTC GCTCGAC); *g53bp1* (*human*) (TCTAGTGTGTTAGATCAGG); *gLin37* (*human*) (TCTAGGGAGCGTCTGGATG).

### Cell cycle phase purification by PIP-FUCCI

Abl pre-B cells or MCF10A cells were transduced with pLenti-CMV-Blast-PIP-FUCCI and selected in 5mg/ml Blasticidin for 3 days(Grant et al. 2018). To collect G_1_ phase cells from proliferating cultures, mVenus-positive cells that were also mCherry-negative were sorted. To collect S/G_2_/M cells, all mCherry-positive cells were sorted. Cell sorting was conducted on a BD FACSAria (BD Biosciences) at the Comprehensive Flow Cytometry Core at University of Alabama at Birmingham (supported by NIH P30 AR048311 and NIH P30 AI27667).

---

**Figure S1.**
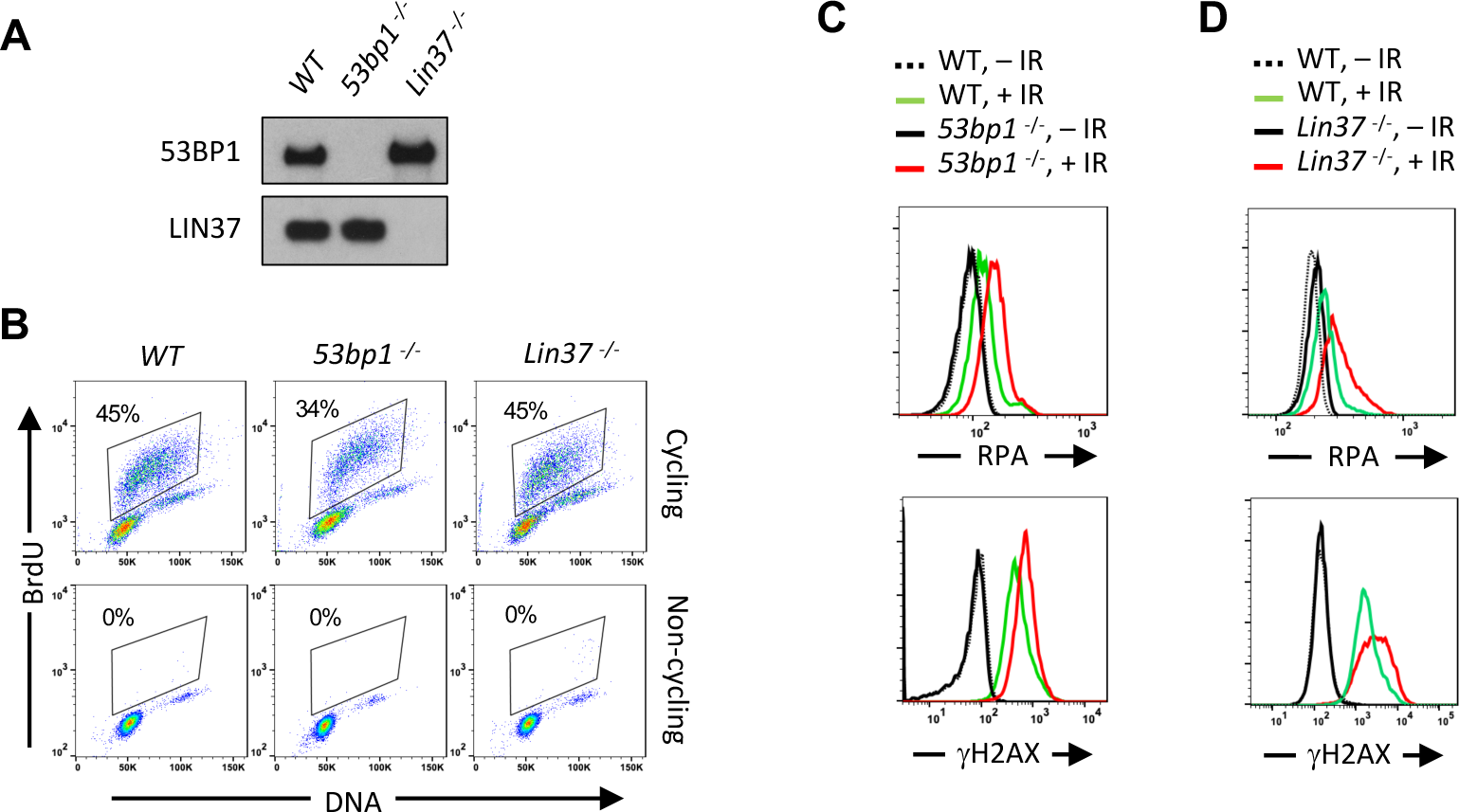
Non-cycling LIN37-deficient cells accumulate chromatin-bound RPA after IR-induced damage (A) Western blot analysis of indicated proteins in *WT*, *53bp1*^-/-^ and *Lin37*^-/-^ abl pre-B cells. (B) Flow cytometric analysis or BrdU and DNA (7AAD) of BrdU pulsed cycling (top) or non-cycling (bottom) *WT*, *53bp1*^-/-^ and *Lin37*^-/-^ abl pre-B cells. Percentage of cells in S-phase is indicated. (C, D) Flow cytometric analysis of chromatin-bound RPA (top panels) and γH2AX (bottom panels) before or after IR of non-cycling *WT* and *53bp1*^-/-^ (C) or *Lin37*^-/-^ (D) abl pre-B cells. The experiments were repeated in two independently generated cell lines at least twice.

**Figure S2.**
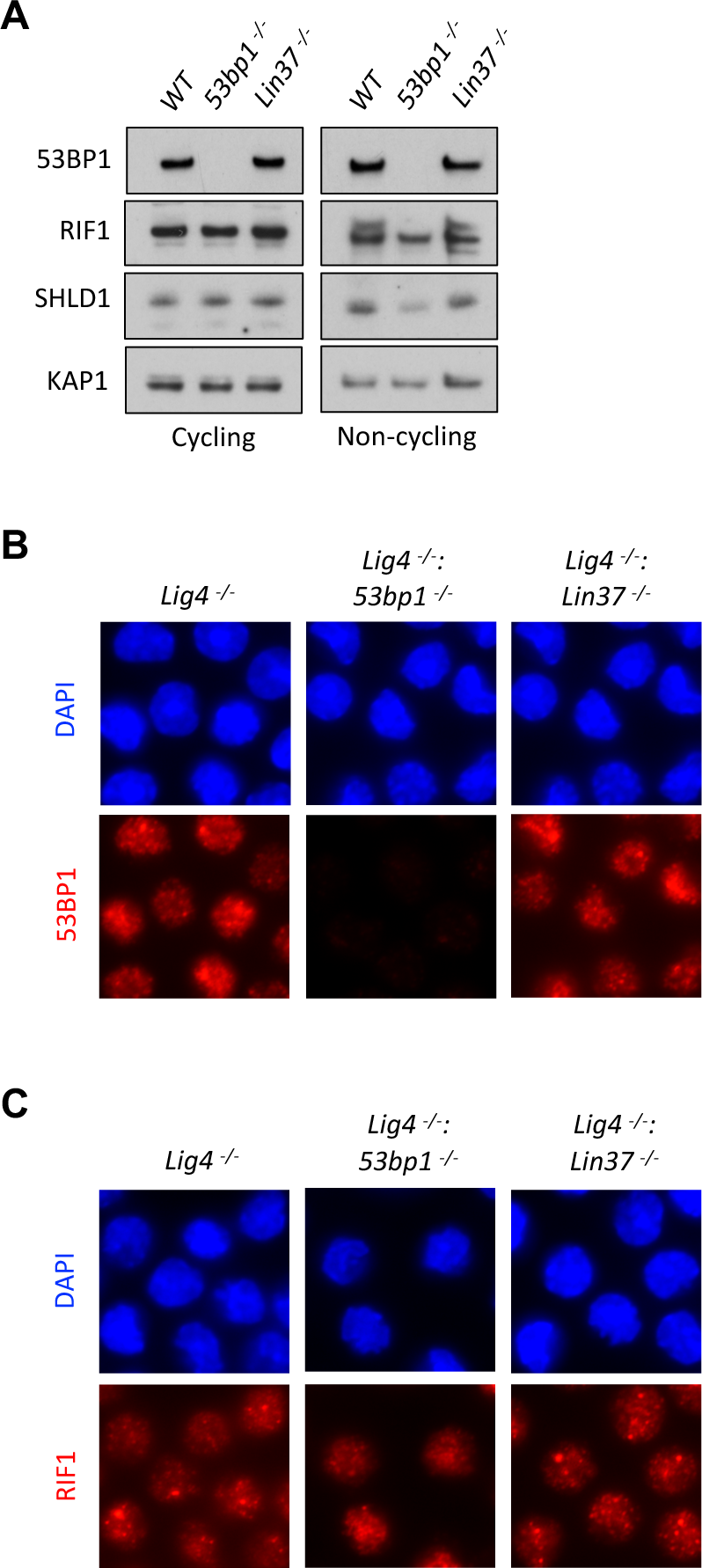
53BP1 and LIN37 have distinct DNA endprotection functions (A) Western blot of indicted proteins in cycling and non-cycling *WT*, *53bp1*^-/-^ and *Lin37*^-/-^ abl pre-B cells. (B, C) Representative images of 53BP1 (B) or RIF1 (C) foci after IR treatment of non-cycling *Lig4*^-/-^, *Lig4*^-/-^:*53bp1*^-/-^ and *Lig4*^-/-^:*Lin37*^-/-^ abl pre-B cells.

**Figure S3.**
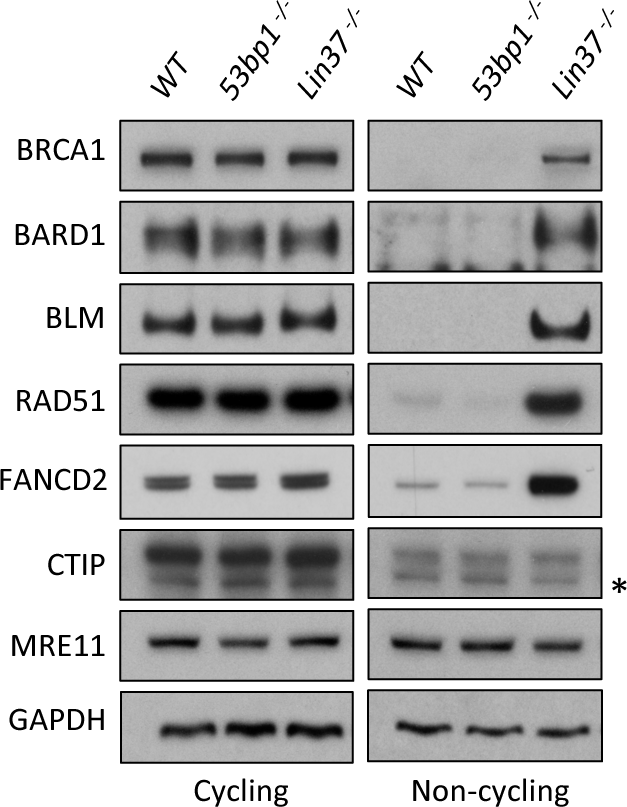
LIN37 suppresses HR protein expression in non-cycling cells. Western blot analysis of indicated proteins in cycling or non-cycling *WT, 53bp1*^-/-^ and *Lin37*^-/-^ abl pre-B cells. * indicates non-specific bands.

**Figure S4.**
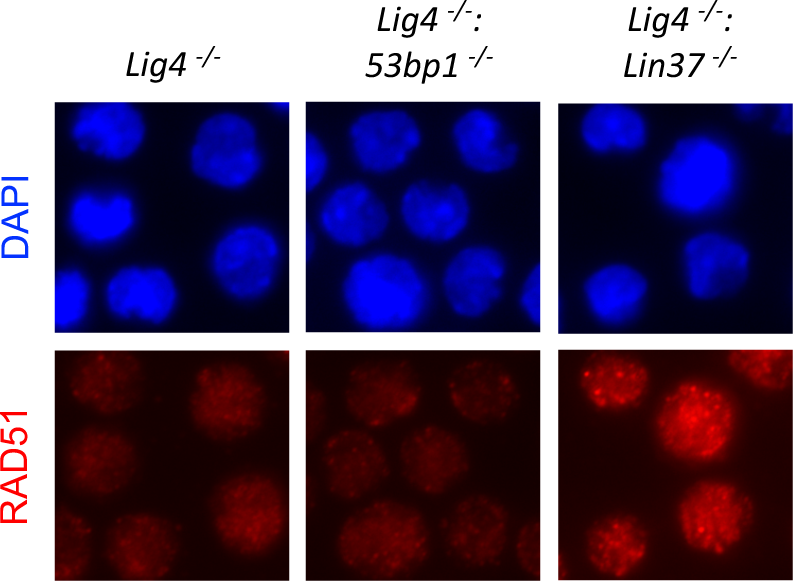
LIN37 deficiency leads to RAD51 focus formation in non-cycling abl pre-B cells Representative images of RAD51 IR-induced foci in non-cycling *Lig4*^-/-^, *Lig4*^-/-^:*53bp1*^-/-^ and *Lig4*^-/-^:*Lin37*^-/-^ abl pre-B cells.

**Figure S5.**
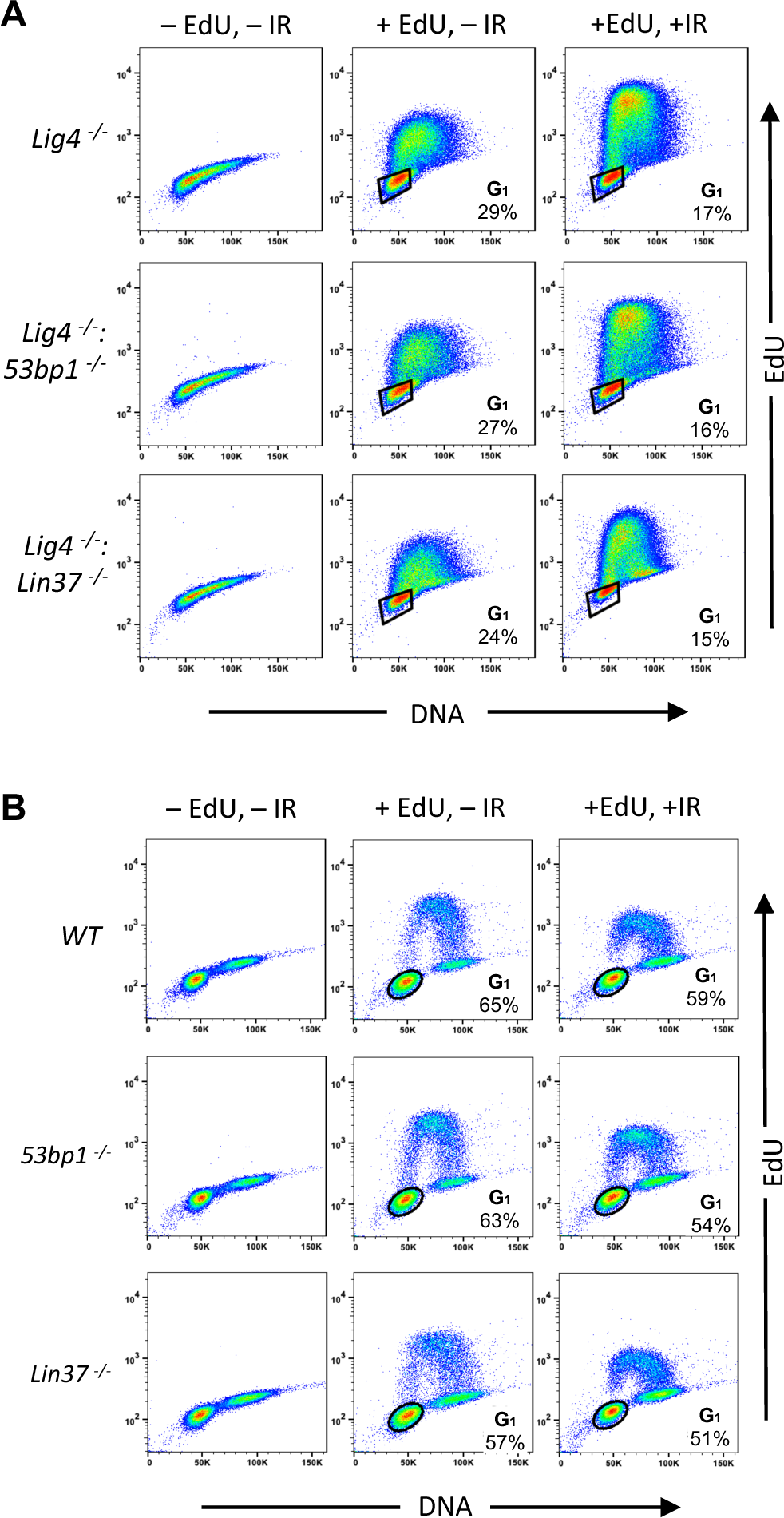
Identification of G_1_ phase cells in proliferating cells (A, B) Flow cytometric analysis of EdU content and DNA (7AAD) of cycling *Lig4^-/-^*, *Lig4^-/-^:53bp1*^-/-^ and *Lig4^-/-^:Lin37*^-/-^ abl pre-B cells (A) or cycling *WT*, *53bp1*^-/-^ and *Lin37*^-/-^ MCF10A cells (B). Cells were treated (+EdU) or not treated (-EdU) with EdU and analyses were carried out before or after IR. The percentage of G_1_ phase cells is shown.

